# An optimal kernel-based method for gene set association analysis

**DOI:** 10.1101/304055

**Authors:** Tao He, Shaoyu Li, Ping-Shou Zhong, Yuehua Cui

## Abstract

Single-variant based genome-wide association studies have successfully detected many genetic variants that are associated with many complex traits. However, their power is limited due to weak marginal signals and ignoring potential complex interactions among genetic variants. Set-based strategy was proposed to provide a remedy where multiple genetic variants in a given set (e.g., gene or pathway) are jointly evaluated, so that the systematic effect of the set is considered. Among many, the kernel-based testing (KBT) framework is one of the most popular and powerful methods in set-based association studies. Given a set of candidate kernels, method has been proposed to choose the one with the smallest p-value. Such a method, however, can yield inflated type I error, especially when the number of variants in a set is large. Alternatively one can get p-values by permutations which, however, could be very time consuming. In this work, we proposed an efficient testing procedure that can not only control type I error rate but also generate power close to the one obtained under the optimal kernel. Our method is built upon the KBT framework and is based on asymptotic results under a high-dimensional setting. Hence it can efficiently deal with the case where the number of variants in a set is much larger than the sample size. Both simulation and real data analysis demonstrate the advantages of the method compared with its counterparts.

## 1 Introduction

Driven by the advancements in microarray and next generation sequencing technologies, increasing number of genetic variants such as single nucleotide polymorphisms (SNPs), indels and copy number variation, are generated in a daily basis. Traditional genome-wide association studies (GWAS), aiming at associating single SNPs with complex traits, have been proven to be a powerful tool to unveil the genetic architecture of many complex traits. However, the power of traditional GWAS analyses by assessing the effect of SNPs one at a time, is limited due to weak marginal signals and the lack of considering potential interactions among genetic variants. Such limitation has been partially addressed by the recent wave of set-based association studies (e.g., Subramanian et al., 2005). The extension to a set-based analysis is a natural choice because genetic variants in a set (e.g., a gene or a pathway) tend to work coordinately to fulfill their joint task. On one hand, the subtle effects in multiple variants can be combined so that the joint signal of the set could be potentially boosted and be detected. On the other hand, the set-based strategy improves the power by capturing the complicated interactions among variants if any. There are a variety of biologically meaningful methods to create a SNP-set or gene-set, such as the annotated gene models (for SNP-set), Kyoto Encyclopedia of Genes and Genomes (KEGG) pathway (Kanehisa, 2000), Reactome (Croft et al., 2011) and Gene Ontology (Ashburner et al., 2000).

The kernel-based testing (KBT) framework, which measures the similarity between genetic variants through a kernel function then compares with the phenotype similarity, is one of the most popular and powerful methods in set-based association studies (Liu et al. 2007; Liu et al. 2008; Kwee et al. 2008). KBT is a very general framework and many other similarity based approaches (e.g., Reiss et al. 2010; Wessel and Schork 2006; Mukhopadhyay et al. 2010; Tzeng et al. 2009) are closely related to it. As observed in the literature (e.g., Wessel and Schork 2006; Wu et al. 2010; Lin et al. 2011), the power of KBT generally depends on the choice of the kernel function. Assuming that the relationship between a gene set and a disease phenotype can be described by a function *h*(·), if the true function *h*(·) comes from the function space generated by the specified kernel, then analysis based on the corresponding kernel will ideally achieve the optimal power. However, the underlying genetic function on a phenotypic response, hence the true function *h*(·), is generally unknown in practice. As a result, it is difficult to choose what kernel should be used. Given a set of candidate kernels in the KBT framework, a common practice is to choose the one leading to the smallest *p*-value. This, however, could inflate the type I error rate due to the choice of kernel selection. To overcome this, Wu et al. (2010) proposed a data dependent perturbation method. However, this strategy is over-conservative in a high-dimensional setting in which the number of variants could be much larger than the sample size. Moreover, it needs computationally intensive procedures to evaluate the statistical significance. The computation burden can further hamper its applicability to large scale genomic data.

In a gene-set association analysis, the number of variants (e.g., SNPs), denoted as *p*, is typically larger than the sample size, denoted as *n*, especially in a pathway-based analysis. Such a large *p* small *n* problem brings statistical challenges in developing a set-based testing procedure. Therefore, our interest is to find an efficient kernel testing procedure that can maintain nominal type I error rate while achieving high power in a high-dimensional setting (i.e., *p* > *n*), under the KBT framework. We mainly focus on a high-dimensional setting and assume a set of candidate kernels are given. We propose an effective and efficient testing procedure when multiple candidate kernels are available. We introduce a new test statistic taking the maximum of the test statistics using the standardized kernels across the candidate set under a high-dimensional setting. We demonstrate the performance of the strategy through a real data application and extensive simulation studies under both continuous and discrete variable settings. The simulation studies show that the proposed approach can maintain the nominal type I error rate, and the maximum method enables the power to be close to the one obtained using the best candidate kernel function in a set, while the perturbation method proposed by Wu et al. (2010) suffers from power loss. Our method enriches the literature of kernel based association methods in genetic association studies, and has broad applications in other fields where the interest is to evaluate the joint (potentially nonlinear) effect of a set of variants with a response.

## 2 Statistical methods

### 2.1 The model setup

We assume that *n* independent subjects from a population are observed in a study design. For the *i*th subject, let *Y_i_* be the quantitative measurement; **X**_*i*_ = (*X_i_*_1_, …, *X_ip_*)^*T*^ be a vector of measurements for a gene set, which could be SNP genotypes in a SNP set, or gene expression profiles in a gene expression set; **W**_*i*_ = (*W_i_*_1_, *W*_*i*2_, …, *W_iL_*)*^T^* be a vector of *L*-dim covariates, where *L* is finite and *i* =1, 2, …, *n*. These covariates can be any clinical variables such as age, gender, and smoking status. In this work, we focus our attention on a *p*-dim SNP set or gene expression set, where *p* is assumed to be large and could be larger than the sample size *n*. For SNP genotype values, an additive model is assumed where *X_ij_* is typically coded as 0, 1 and 2 corresponding to the number of minor alleles that subject *i* possesses at the *j*th specific locus. The gene expression values are measured as intensity in microarray studies or FPKM values in RNA-seq studies. In this work, we do not assume any specific distribution assumption on **X**_*i*_. This makes our method more general in which it can deal with gene set based association analysis for both gene expression and SNP data. In the follows, we use gene set to denote a SNP set or gene expression set.

To model the relationship between a quantitative trait and a gene set, we consider the following semi-parametric regression model,

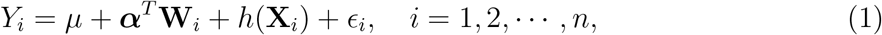

where *h*(·) is an unknown function, *є_i_* is a random subject-specific error term following a certain distribution (not necessarily normal) with E(*є_i_*) = 0, Var(*є_i_*) = *σ*^2^ and is independent of (**X**_*i*_, **W**_*i*_). The identifiability of the *h* function is assured by the side condition E[*h*(**X**_*i*_)] = 0. Our interest is to test association between a gene set and a continuous trait of interest, which can be done by testing the following hypotheses,

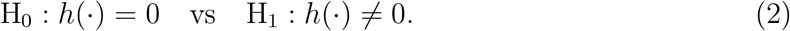

### 2.2 Kernel function

Our method is built upon the KBT idea (Liu et al. 2007; Kwee et al. 2008), but with a different testing strategy. Before proceeding to the KBT statistic, we introduce some basics about the kernel function, which is widely used to measure the similarity between two subjects. Kernel function is commonly used to generate the functional space for the underlying true function *h*(·). A function 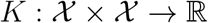 is called a kernel function if it is symmetric and positive semi-definite (i.e., *K*(*x*_1_,*x*_2_) = *K*(*x*_2_,*x*_1_) for any 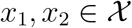, and the *N × N* kernel matrix 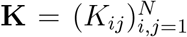 is positive semi-definite, for any 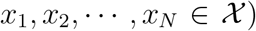 with the (*i*,*j*)-th component *K_ij_* = *K*(*x_i_*,*x_j_*). In our context, *K*(**X**_*i*_, **X**_*j*_) is a measure of similarity between the *i*th and the *j*th subject based on the SNP genotype or gene expression values.

For any positive definite kernel *K*^*^ with corresponding matrix **K**^*^, we can defined its centralized kernel

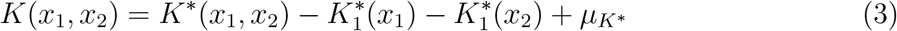

satisfying E{*K*(*X*_1_,*X*_2_)} = 0, where 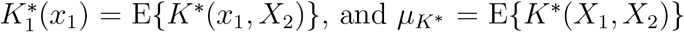. Empirically, centralized kernel matrix **K** can be replaced by its estimator

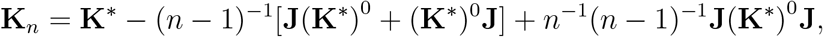

where **J** is an *n* × *n* matrix with all the elements as 1, and **D**^0^ = **D** - diag(**D**) is a zero-diagonal matrix defined for any square matrix **D** which sharing all non-diagonal elements with **D**. For notation simplicity, hereafter we use *K^*^, K* and **K**_*n*_ to represent the original kernel function, centralized kernel function and the empirical version of the centralized kernel matrix, respectively.

Some commonly used kernel functions include linear kernel 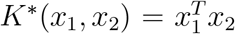, polynomial kernel 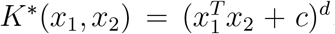, Gaussian kernel 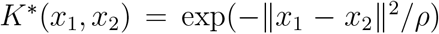 where *c*, 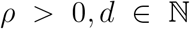 are tuning parameters, and IBS kernel defined as 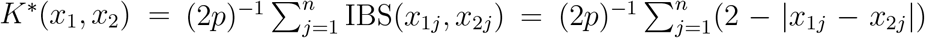. The IBS kernel is for discrete genotype data only. For a review of genomic similarity and more kernel functions, please refer to Schaid (2010a, 2010b).

Throughout this work, we focus on centralized kernel in the testing since the asymptotic distribution of the test statistic using non-centralized kernel is largely determined by the centralized kernel except a location shift. More benefits of using centralized kernel can be found in Lindsay et al. (2008, 2014). Furthermore, we can define the standardized kernel

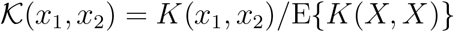

from which it is easy to verify that 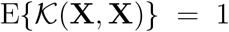. Next let us briefly look at the eigen-decomposition of a kernel function, which is an important way to characterize a kernel function. Assume *K*(·, ·) is a kernel function defined on 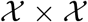. Then the spectral decomposition theorems (Lemma 1 of Chapter 2, Steinwart and Scovel, 2012) implies that the standardized kernel 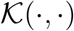 enjoys the following representation

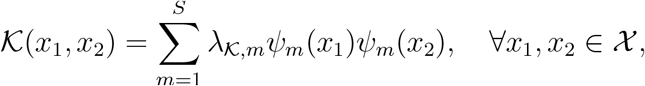

where the eigenfunctions 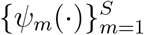 form a complete orthonormal system (i.e., 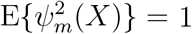 for any *m*, 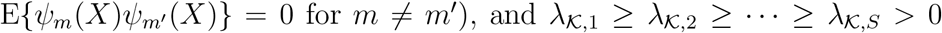 are the non-zero eigenvalues satisfying 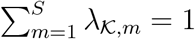. The standardization is required because E{*K*(**X**, **X**)} could diverge in the high-dimensional case, and it ensures 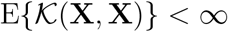 so that the eigen-decomposition can be properly defined. By denoting 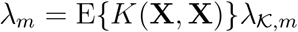, we can get the pseudo eigen-decomposition of kernel function *K*(·, ·)

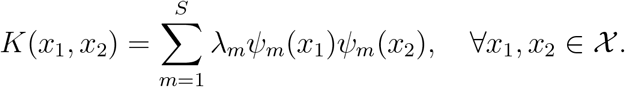

It should be noticed that the eigen-decomposition not only depends on the expression of the kernel, but also implicitly depends on the space 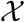 (e.g., dimension *p*).

A functional space 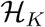, namely a reproducing kernel Hilbert space (RKHS), can be generated by any positive semi-definite kernel function *K*(·, ·). The form of the functions that reside in 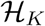 is characterized by the kernel function *K*. Here we assume that the *h*(·) function in model (1) is a member of the RKHS 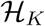. Therefore, by specifying the kernel function, we assume that *h*(·) function has some structure defined by 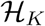. For example, linear kernel indicates that the overall genetic effect is a linear combination of the individual effects in the set, i.e., *h*(**X**_*i*_) = *β^T^***X**_*i*_; polynomial kernel with (*c, d*) = (1, 2) implies a quadratic model 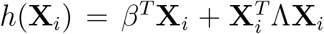, where interactions are modeled in addition to the linear effects, *β* and Λ are coefficient vector and matrix, respectively. Because different kernel functions are associated with different functional spaces, the kernel based approach is very flexible for modeling different types of functions as well as complicated (potentially nonlinear) interactions among variants. On the other hand, challenges arises given that the true function is generally unknown in practice. It is expected that the power of KBT is limited if a kernel function is misspecified. In the following sections, we start with the hypothesis testing problem using a single kernel function, followed by the ones using multiple kernel functions through which the power can be greatly boosted.

#### 2.3 Hypothesis test based on a single kernel

We consider the following kernel-based U-statistic (KU)

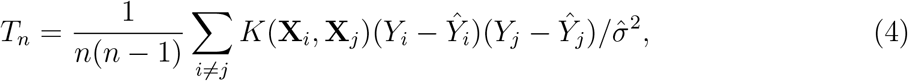

where 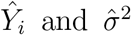 are the sample estimates under the null model 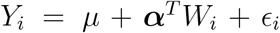. Specifically, let 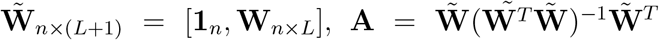, then 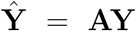 and 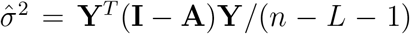. Define 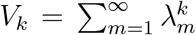 for any positive integer *k*. Then the asymptotic normality of the test statistic *T_n_* under the null hypothesis is stated in the following theorem.

##### Theorem 1

*Assume the density function of error ∊ is symmetric around 0 with* 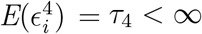. *Then, (i) under the null hypothesis of no genetic effect (i.e., h*(·) = 0),

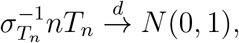

*if*

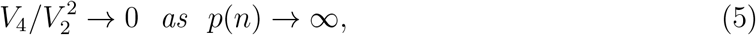

*where* 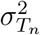 *is the variance of nT_n_ and can be estimated by the following estimator*

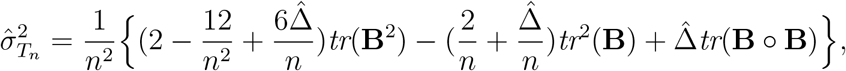

*where* 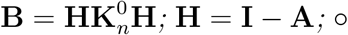 *denotes the Hadamard product (elementwise product); and* 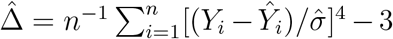. *(ii) Assume that* 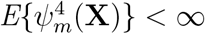 *for all integers *m*. Under the local alternative H*_1*n*_ : *h*(x) = *d_n_*(x), *where d_n_ satisfies two conditions:* 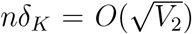 *with* 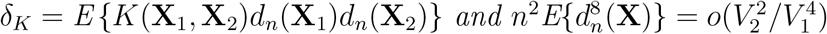, *we have*

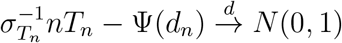

*if (5) holds, where* 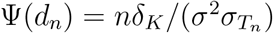 *is the location shift*.

A sketch of proof to Theorem 1 is relegated to Appendix. Given the asymptotic normality, we can then obtain the *p*-value for testing *H*_0_ : *h*(.) = 0, i.e.,

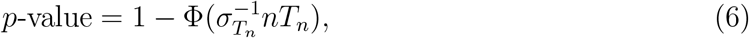

where Φ(·) is the cumulative density function for a standard normal distribution. As we can see from Theorem 1, the asymptotic normality holds if the condition 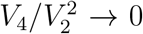 holds. It was mentioned earlier that this ratio depends on the kernel function, the dimension *p* of the space where the kernel is defined, and the probability measure on 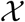. To highlight the effect of dimension, define 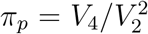. In the following, we take a further look at the conditions for some commonly used kernel functions.

##### Example 1

*Consider the linear kernel* 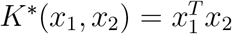, *and assume a multivariate random variable* **X**_*i*_ = (*X_i_*_1_, …, *X_ip_*) *with covariance matrix* Σ, *i* = 1, …, *n. Then π_p_* = *tr*(Σ^4^)/*tr*^2^(Σ^2^).

##### Example 2

*Consider the quadratic kernel* 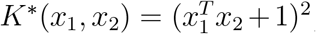, *which is a special polynomial kernel. Denote* 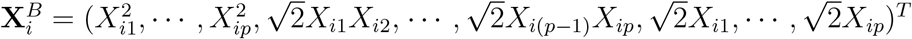 *as a J-dim random vector with covariance matrix* Σ_B_, *where J* = (*p*^2^ + 3*p*)/2. *Then* 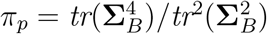.

##### Example 3

*Consider the IBS kernel*

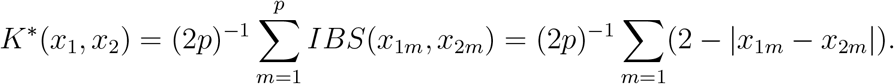

*Denote* 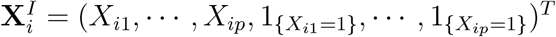 *as a* 2*p-dim random vector with covariance matrix* Σ_*i*_. *Then* 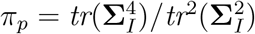.

The proofs to Example 1–3 are relegated to Appendix. From the above examples we can see that under the three widely-used kernels, condition (5) is equivalent to a condition on the covariance matrix of a random vector whose length depends on *p*. Besides, it is a weak condition that brings little constraint to the growth rate of *p* relative to *n*. Moreover, if all the eigenvalues of the covariance matrix Σ is bounded, then it is not difficult to see that *π_p_* is of orders *p*^−1^, *p*^−2^ and *p*^−1^ respectively for the linear, quadratic and IBS kernel defined earlier, and *π_p_* ⟶ 0 as *p* ⟶ 0. For more discussion on the condition tr(Σ^4^)/tr^2^(Σ^2^) ⟶ 0, please refer to Chen et al. (2010).

Although the explicit condition for the covariance matrix of many kernel functions is typically unknown, there do exist consistent estimators for *V*_2_ and *V*_4_ that can provide us the empirical version of *π_p_*. Specifically, 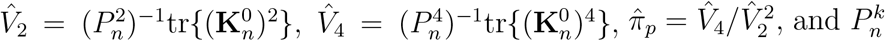 is the number of *k*-permutations of *n*.

### 2.4 Hypothesis test under multiple candidate kernels

In the previous section, we proposed a test statistic based on a single candidate kernel, and we showed its asymptotic normality under a high-dimensional setting. Since the optimal kernel is generally unknown in practice, we consider a set of *M* (finite) candidate kernel functions *K*_1_(·, ·), *K*_2_(·, ·), …, *K_M_* (·, ·) with kernel matrix **K**_*n*,1_, **K**_*n*,2_, ···, **K**_*n,M*_. Two testing methods are proposed under this setting. In the first one, a new kernel function is generated by taking the simple average of the normalized candidate kernels and then apply it to the single kernel based testing procedure. The second method uses a maximum test statistic and the well-developed results on multivariate normal distribution. Both methods are computationally efficient and easy to implement in practice.

#### 2.4.1 Test based on kernel average

Without any prior knowledge of the nonparametric function *h*(·) in (1), taking the simple average among a set of normalized kernels is a natural choice, where the normalization is necessary for equal-metric consideration. In particular, denote the standardized kernels with their empirical matrix forms as

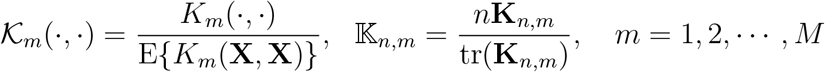

and the simple average kernel with its matrix form as

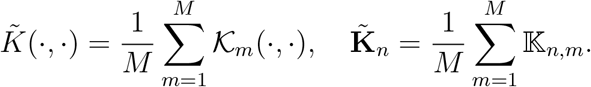

Intuitively, the performance of the test using 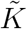 is most likely a compromise between the best and the worst ones. Its power will not be close to the optimal one among a candidate set, but it is a conservative option to improve the power over the weakest choice in the set given the fact that the truth is unknown in practice. We call this test as the simple average test.

#### 2.4.2 Maximum test among a candidate set

An alternative strategy to the average kernel testing is to perform the test for individual kernels, then taking the maximum as the test statistic. Taking the maximum test statistic is the same as taking the minimum *p*-value which has been proposed in literature. However, the minimum *p*-value method often requires computationally expensive techniques such as permutation or perturbation to evaluate the null distribution. Here we focus on the maximum test statistic among all the candidate kernels and take advantage of the derived asymptotic normality under the high dimensional assumption. Let *nT_n,m_* and 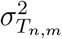 be respectively the test statistic and the corresponding variance using the *m*th kernel function, and denote 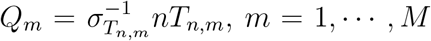. As we can see from (6), the *p*-value is fully determined by *Q_m_*, hence maximizing *Q_m_* is equivalent to minimizing a nonlinear function of *p*-values. We focus on the following maximum statistic

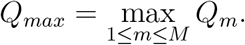

Let *ρ_kl,n_* = cov(*Q_k_, Q_l_*) and 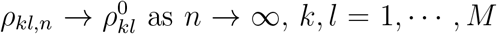. The following theorem states the the asymptotic distribution of the maximum statistic *Q_max_*.

##### Theorem 2

*Assume condition (5) in Theorem 1 is satisfied for each candidate kernel K_m_, then*

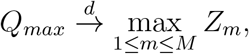

*where* **Z** = (*Z*_1_, *Z*_2_, …, *Z_M_*)*^T^ follows a multivariate normal distribution with mean* **0**_*M*_ *and covariance matrix* 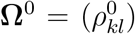. *Moreover, under the local alternative H*_1*n*_, *the power of the maximum test achieves what the optimal one does among a candidate set, when the location shift of the optimal kernel, specified in Theorem 1, is large enough*.

The proof of Theorem 2 is relegated to Appendix. Based on Theorem 2, the *p*-value of the maximum test can be calculated as

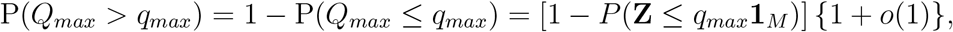

where the leading order term can be efficiently and accurately calculated in many popular platforms (e.g., *mvnorm* package in R). Although the true covariance matrix **Ω**^0^ is unknown, it can be approximately substituted by its consistent estimator 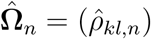, where

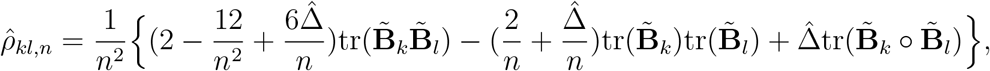

where 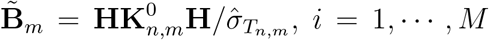. This maximum test strategy enjoys several merits. First, the nature of maximum strategy enables the best power among a set of candidate kernels. Second, the asymptotic normality results obtained under the high-dimensional asymptotics greatly reduce our computational burden, and protects the size from being inflated or over-conservative. Although the maximum method is designed for the high-dimensional case, we found in the extensive simulation studies that the method is also applicable when the dimension *p* is low. Specifically, type I error rate was well-protected and only slightly conservative when *p* is very low (e.g., *p* = 10). Under a low-dimensional case, the distribution of *Q_max_* can be approximately viewed as the maximum among *M* correlated chi-square random variables. Although its asymptotic behavior is beyond the scope of this paper, from our simulation studies, we found that as *p* grows (*p* ≥ 20), the empirical type I error is very close to the nominated level and the shape of the distributions *Q_m_*(*m* = 1,…, *M*) gets closer to a normal distribution. As the number of variants in a gene set (e.g., in a pathway) is typically large, the proposed test is generally safe to apply in practice. We call this test as the maximum test.

## 3 Simulation studies

Extensive simulation studies were conducted to evaluate the type I error rate and the empirical power of the proposed methods. A continuous trait was simulated from the following model,

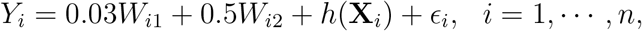

where *∊_i_* are independent and identically distributed random errors generated from *N*(0,1) distribution, *W*_*i*1_ ∼ *N*(2,1) and *W*_*i*2_ ∼ *Ber*(0.6) are independent covariates, and **X**_*i*_ is a *p*-dim discrete or continuous vector representing genotypes or gene expression profiles. To evaluate the type I error, we generated data sets under the null hypothesis of no association (i.e., *h*(·) = 0), and recorded the proportion of (incorrectly) rejecting the null hypothesis. To assess the power, we generated data sets by specifying the *h* function, and recorded the proportion of (correctly) rejecting the null hypothesis. We conducted 1000 simulation replications in each case and set the significance level as 0.05. In the following, we assessed the performance of the proposed methods under the continuous and discrete variant settings separately.

### 3.1 Continuous variants

Under the continuous variant setting, we simulated **X**_*i*_ = (*X_i_*_1_, …, *X_ip_*) from a multivariate normal distribution with mean **0**_*p*_ and covariance matrix 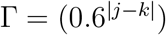, where *p* = 50,100 and *i* =1, …, *n*. The sample was assumed to be *n* = 500,1000, 2000. The candidate set consists of three commonly used kernels, including linear kernel, polynomial kernel (*c* = 1, *d* = 2) and Gaussian kernel 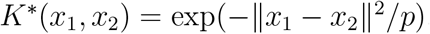. In addition to a single kernel based test, the kernel average method (denoted as SimplyAv), the perturbation method (denoted as Pertb) (Wu et al. 2010) and the maximum method (denoted as Max) were also applied. Table 1 reports the type I error rates of tests with varying sample size. We can see that the type I error was not well-protected using the perturbation method, and others are reasonably controlled (close to the nominal level 0.05). This finding implies that the perturbation method is relatively conservative under the high-dimensional setup, while the other method works reasonably well.

**Table 1:** Empirical type I error rates of different tests under the continuous variant setting

| $p$ | $n$ | Gau | Linear | Poly | SimplyAv | Pertb | Max |
| --- | --- | --- | --- | --- | --- | --- | --- |
| 50 | 500 | 0.061 | 0.056 | 0.046 | 0.056 | 0.020 | 0.054 |
|  | 1000 | 0.055 | 0.058 | 0.047 | 0.057 | 0.025 | 0.056 |
|  | 2000 | 0.052 | 0.048 | 0.050 | 0.050 | 0.036 | 0.049 |
| 100 | 500 | 0.055 | 0.051 | 0.063 | 0.051 | 0.012 | 0.058 |
|  | 1000 | 0.055 | 0.055 | 0.056 | 0.053 | 0.015 | 0.058 |
|  | 2000 | 0.052 | 0.051 | 0.046 | 0.051 | 0.021 | 0.047 |

To evaluate the testing power, we considered four different scenarios. Under each scenario, the *h*(·) function was set differently as follows:

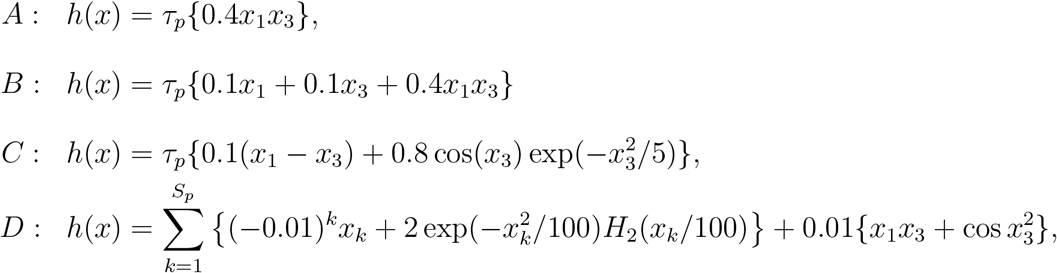

where *H_k_*(·) is the *k*th order Hermite polynomial, *τ_p_* and *S_p_* are two constants that were set differently for each *p* to adjust for the overall effect. Specifically, (*τ_p_, S_p_*) = (0.8,8) when *p* = 50, and (*τ_p_, S_p_*) = (1, 30) when *p* = 100. For each scenario, 1000 simulation replicates were generated to estimate the empirical power. Figure 1 and Figure 2 show the empirical power under different scenarios for *p* = 50 and *p* = 100 respectively. We can see that different kernels have different powers, depending on the underlying trait architecture. Simple average kernel gives intermediate power among the candidate kernels, and the power of maximum test under each scenario was generally close to the optimal kernel. For example, under scenario A the polynomial kernel was the optimal kernel in terms of best power. Among the three competitive ones (i.e., SimplyAv, Pertb and Max), the maximum test gives power more close to the best one and performs the best among the three. The same pattern can be seen under other three scenarios for both *p* = 50 and *p* =100 settings. It is also worth mentioning that the perturbation method suffers tremendously from power loss when the sample size is small (see cases with *n* = 500 and *n* = 1000). This implies that the Pertb method was not suitable for large *p*. The simulation study demonstrates that the maximum strategy is a good solution in practice to maintain proper power over the weak choices of kernels under the high-dimensional setting.

**Figure 1:**
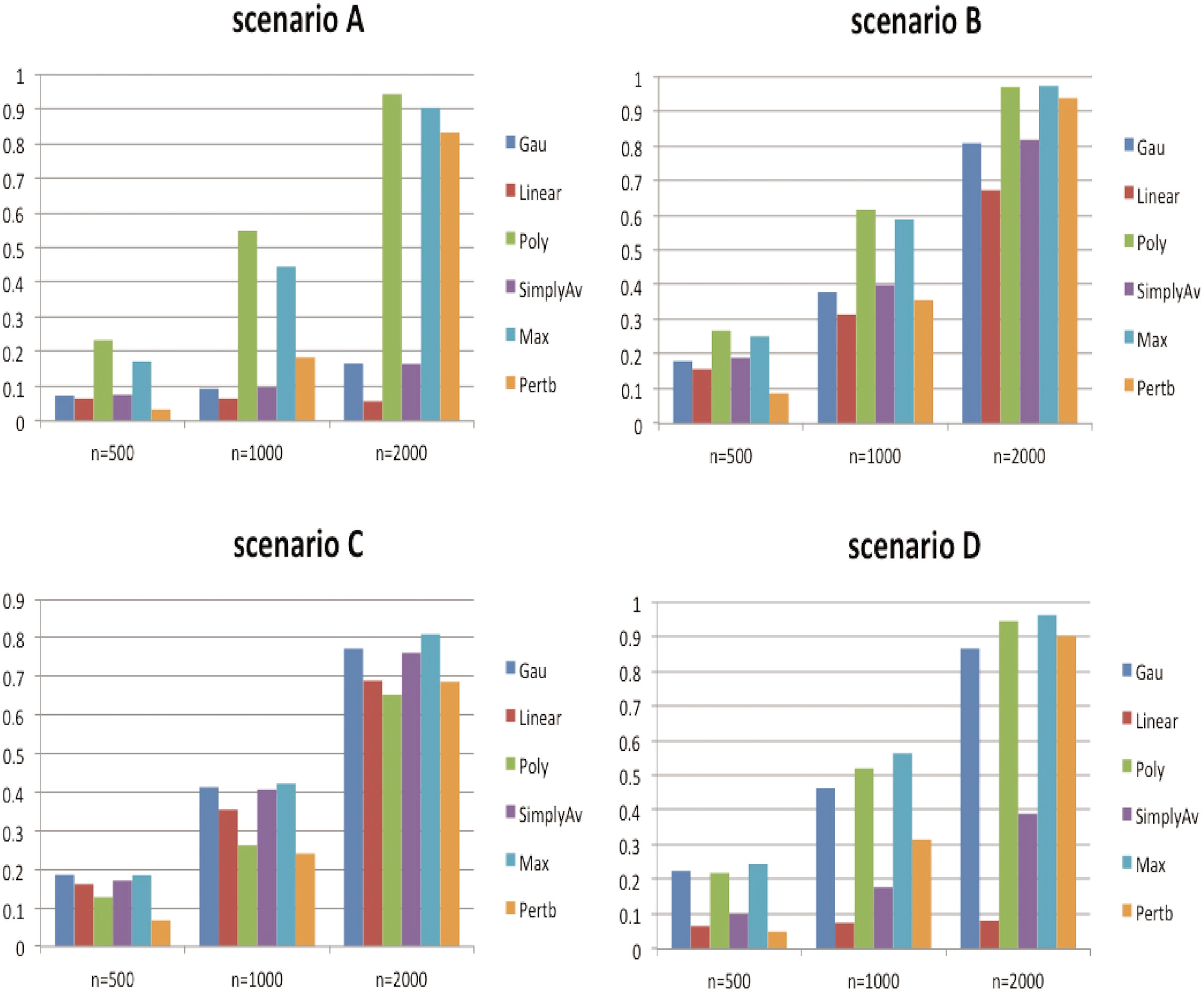
Empirical testing power of different tests under different scenarios and sample sizes with the continuous variant setting when *p* = 50.

**Figure 2:**
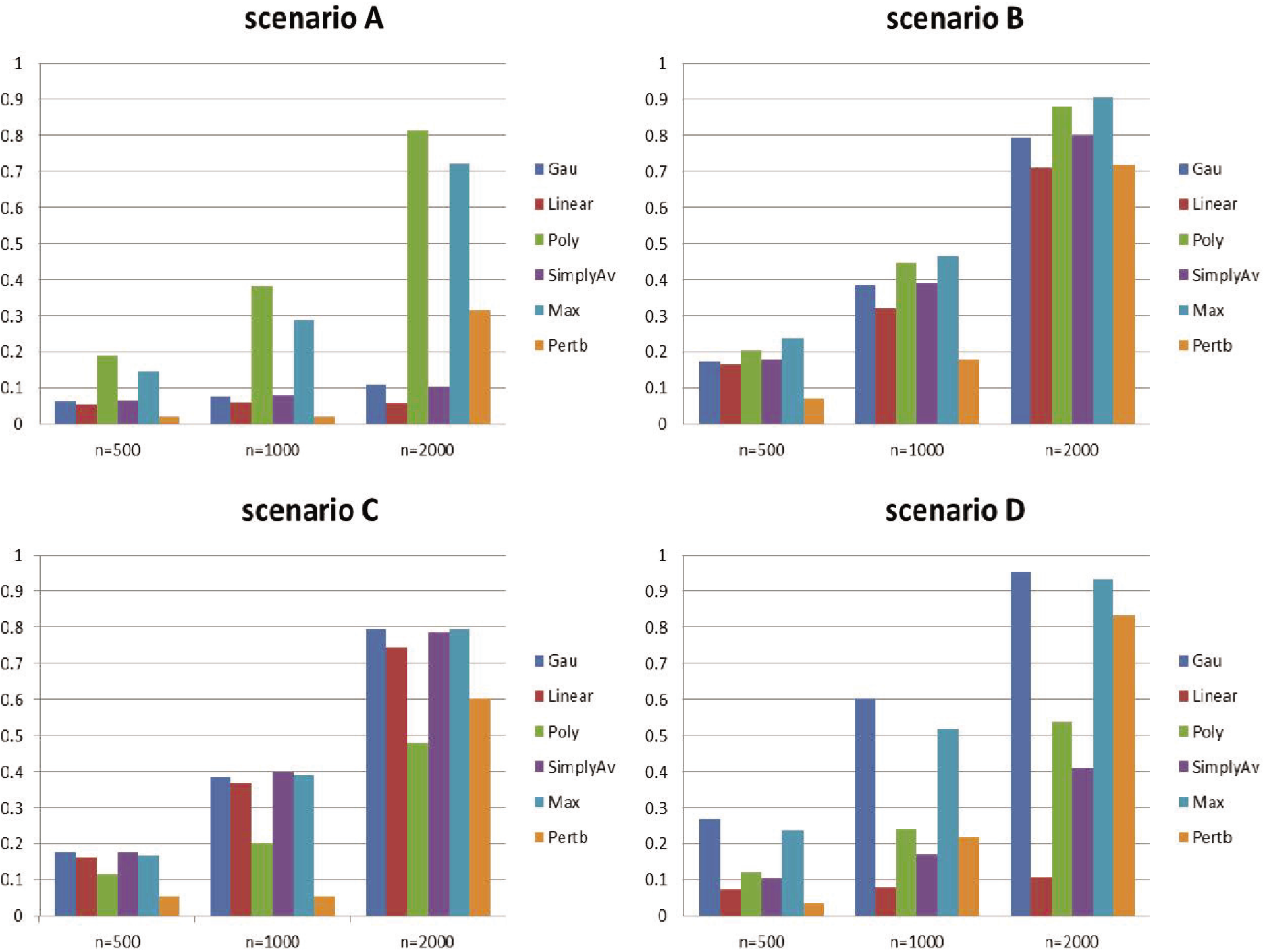
Empirical testing power of different tests under different scenarios and sample sizes with the continuous variant setting when *p* = 100.

### 3.2 Discrete variants

For the discrete variant setting, we generated genotypes based on 378 HAPMAP SNPs located within the KEGG thyroid cancer pathway using the HAPGEN software (Marchini et al. 2007). This pathway was detected as a significant pathway associated with birth weight in our real data analysis given in Section 4. We simulated the quantitative trait for *n* = 1000, 2000, under three scenarios E, F, and G. Under scenario E, we let the *h*(·) function take the form of

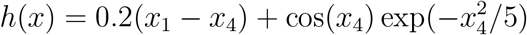

where the fourth SNP has a nonlinear effect on the response in addition to the main effects of SNP 1 and 4 with different effect directions.

To mimic the situation where a large number of SNPs contributes to the trait variation, we considered the following model,

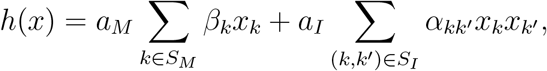

where *S_M_* is a pre-defined set of 30 SNPs with main effects, *S_I_* consists of 60 SNP-pairs representing 60 simple interactions. Both {*β_k_,k* ∈ *S_M_*} and {*α_kk’_* (*k,k’*) ∈ *S_I_*} were independently generated from *Uniƒ* (0, 0.02), and were fixed once generated for all simulation replicates. We set the coefficients (*a_M_, a_I_*) = (0.01,1.5) under scenario F, indicating the combination of weak main effects and relatively strong interaction effects. We let (*a_M_, a_I_*) = (3.5, 0) under scenario G, which implies a pure main-effect model.

In addition to linear and polynomial kernels, we added the IBS kernel to the candidate set, since it is commonly used to measure SNP similarity between two subjects in genetic association studies. Similar to the previous section, the SimplyAv, Pertb and Max methods were applied. Table 2 displays the type I error rates of different tests under different sample sizes. We can see that all the tests maintained reasonable type I error rate except the Pertb method which is a little conservative. Again, the reason might be due to the high dimensionality of which the Pertb method cannot handle very well.

**Table 2:** Empirical type I error rates of different tests under the discrete variant setting.

| $n$ | IBS | Linear | Poly | SimplyAv | Pertb | Max |
| --- | --- | --- | --- | --- | --- | --- |
| 1000 | 0.052 | 0.050 | 0.047 | 0.050 | 0.037 | 0.054 |
| 2000 | 0.053 | 0.045 | 0.046 | 0.043 | 0.038 | 0.050 |

The power simulation results are shown in Table 3, where the best and second best powers among all the tests are shown with the underline and bold font, respectively. Again, we observed the power difference of applying different kernels. Among the different methods, the perturbation method has the smallest power which might be due to the issue of high-dimensionality. The maximum test always achieves the power as close as the best power indicating the robustness of the testing procedure by taking the maximum among the three individual ones. We also noticed the power improvement as the sample size increases.

**Table 3:** Empirical power of testing with single kernel and multiple kernels under the discrete variants setting^*^

| $n$ | Scenario | IBS | Linear | Poly | SimplyAv | Pertb | Max |
| --- | --- | --- | --- | --- | --- | --- | --- |
| 1000 | E | <u>0.526</u> | 0.457 | 0.429 | 0.480 | 0.388 | **0.488** |
|  | F | 0.397 | 0.412 | <u>0.475</u> | 0.428 | 0.383 | **0.452** |
|  | G | 0.390 | 0.423 | <u>0.444</u> | 0.422 | 0.356 | **0.431** |
| 2000 | E | <u>0.967</u> | 0.932 | 0.913 | 0.954 | 0.927 | **0.961** |
|  | F | 0.738 | 0.753 | <u>0.813</u> | 0.781 | 0.745 | **0.796** |
|  | G | 0.769 | 0.790 | <u>0.808</u> | 0.798 | 0.748 | **0.799** |

\* The best power across all the tests is underlined, and the second best is shown as bold font.

In summary, the simulation results indicate that it is generally safe to apply the maximum test strategy given a set of candidate kernels. The maximum test can control the type I error reasonably well, while it also maintains relatively high power. Without knowing the underlying truth, the maximum test procedure is safely recommended in practice under a high-dimensional setup.

## 4 Application to real data

We illustrated our methods via the analysis of a Thai baby birth weight data set to investigate significant pathways that are associated with birth weight. As part of Hyperglycemia and Adverse Pregnacy Outcome (HAPO) study, this data collect genotype and phenotype information for 1209 Thai infants and their mothers. For more details about the HAPO study, please refer to http://www.ncbi.nlm.nih.gov/projects/gap/cgi-bin/study.cgi?study_id=phs000096.v4.pl&phv=163690&phd=2831&pha=&pht=2446&phvf=&phdf=&phaf=&phtf=&dssp=1&consent=&temp=1. We removed infants with large proportion of missing SNPs (> 10%), and SNPs with minor allele frequency (MAF) less than 0.05 or showing deviation from Hardy-Weinberg equilibrium (*p*-value< 0.001). The final data set contains 970,342 SNPs in 1189 infants (580 males, 509 females). The pathways were defined by Kyoto Encyclopedia of Genes and Genomes (KEGG) (Kanehisa and Goto, 2000). SNPs that are within 5kb up- and down-stream of a gene were firstly assigned to the corresponding gene based on Human Genome Build v38, and then grouped into 186 pathways based on the KEGG pathway information retrieved from the Molecular Signature Database (MSigDB) (Subramanian et al., 2005). The size of the pathways ranges from 167 to 9,912 (SNPs), where > 86% of the pathways are of dimension higher than 500.

We tested the association of each pathway with birth weight, adapting gender (1=male, 2=female) and baby’s gestational age at delivery (in weeks) as two covariates. Since we had little knowledge about the underlying true functional mechanism, we applied three different kernels in the test, including IBS kernel, linear kernel and polynomial kernel (*c* = 1, *d* = 2). We applied simple average kernel test, the perturbation method (Wu et al. 2010) and the maximum statistic method. The false discovery rate was controlled using *q*-value significance levels (0.05 and 0.1) (Storey and Tibshirani, 2003). Table 4 summarizes the significant KEGG pathway index using different methods. The corresponding *p*-values and information of the significant pathways are reported in Table 5.

**Table 4:** Significant KEGG pathway index using different methods.

| $q$-level | IBS | Linear | Poly | SimAv | Perturb | Max |
| --- | --- | --- | --- | --- | --- | --- |
| 0.10 | {36,44,101,169} | {36,80,123,169} | {36,48,80,169} | {36,80,169} | NA | {36,44,80,101,169} |
| 0.05 | {101,169} | {36,80,169} | {36,169} | {169} | NA | {36,101,169} |

**Table 5:** List of significant KEGG pathways and the *p*-values using the corresponding kernel functions.

| idx | # of SNPs | Name* | IBS | Linear | Poly | SimAv | Max |
| --- | --- | --- | --- | --- | --- | --- | --- |
| 169 | 485 | KTC | 1.93E-05 | 1.42E-07 | 1.32E-08 | 1.95E-07 | 1.32E-08 |
| 101 | 914 | KP | 1.71E-04 | 2.27E-02 | 2.38E-02 | 5.50E-03 | 2.84E-04 |
| 36 | 785 | KGBCS | 3.13E-03 | 8.14E-04 | 5.25E-04 | 9.76E-04 | 8.54E-04 |
| 80 | 914 | KPSP | 1.16E-02 | 1.06E-03 | 1.53E-03 | 2.20E-03 | 1.77E-03 |
| 44 | 1052 | KAAM | 1.38E-03 | 8.06E-02 | 8.28E-02 | 2.68E-02 | 2.32E-03 |
| 48 | 419 | KGBLANS | 3.55E-02 | 5.18E-03 | 3.19E-03 | 7.56E-03 | 5.41E-03 |
| 123 | 555 | KNLRSP | 1.32E-02 | 3.78E-03 | 7.43E-03 | 6.02E-03 | 5.99E-03 |

\*KTC: KEGG thyroid cancer; KP: KEGG peroxisome; KGBCS: KEGG glycosaminoglycan biosynthesis chondroitin sulfate; KPSP: KEGG ppar signaling pathway; KAAM: KEGG arachidonic acid metabolism; KGBLANS: KEGG glycosphingolipid biosynthesis lacto and neolacto series; KNLRSP: KEGG nod like receptor signaling pathway.

The result shows that the perturbation method (Wu et al. 2010) fails to detect any signal, which is probably due to the over-conservative behavior under the high-dimensional setting. Among the seven distinct pathways detected by the three kernels at *q*-level 0.1, the maximum test was able to capture five of them, while individual kernel and simple average kernel identified four and three of them, respectively. At *q*-level 0.05, the observations were quite similar. One important observation is that the *p*-value of the maximum test is generally close to the smallest *p*-value among the three kernels, which implies that the maximum test tends to improve the power over the weak choice of kernels. Simply taking average did not achieve the power as the maximum test did.

## 5 Discussion

In this work, we developed testing procedures to test relationship between multiple variants in a gene set and a quantitative trait, while adjusting for other covariates’ effects. We considered a general setting where the variants work coordinately in a (non)linear way, and the dimension of the variants *p* is high in the sense that *p* can go to infinity as sample size *n* goes to infinity. We first proposed a test statistic based on a single kernel function, and derived its asymptotic distribution under the null hypothesis. Based on this, we proposed a practical and efficient testing strategy when multiple candidate kernels are available. We demonstrated, via extensive simulation studies and real data analysis, that under a high-dimensional setting the maximum method can reasonably control the false positive rate while they can also improve the power over a set of weaker choices of kernels. In particular, the maximum method performs as good as the optimal one for a given set of candidate kernels, hence should be recommended in practice. Compared to the perturbation method (Wu et al., 2013), the maximum method outperformed it uniformly in various simulation settings.

Our methods enjoy several advantages as described below. The first advantage lies on the ability to accommodate high-dimensional variants and to maintain reasonable type I error rate, even if the utilized kernel functions do not reflect the underlying relationship between the variants and the trait. Another advantage is the flexibility, which is revealed in two aspects. On one hand, we consider a general model which can potentially capture any complex interaction mechanism and is different from many models restricted to linear relation and/or linear interactions. On the other hand, when there are a range of kernels that can be selected to form the candidate set, the proposed maximum kernel testing strategy is shown to maintain improved power over the poor choices of kernels in the set, without the prior knowledge of the underlying genetic function.

Thirdly, our method is easy to implement and is free of computational burden, by applying the asymptotic result of the test statistic. This can greatly facility the applications in pathway (or gene-set) association studies where the variants (SNPs or gene expression profiles) are typically in high dimensions. The unique feature of our method under a high-dimensional setup distinguishes itself from many existing ones. Given the typical norm of high-dimensionality in gene set association studies, our method should be a good choice to implement. Our method relies on the asymptotic results where the dimension *p* is relatively large. Although large *p* is very typical in gene set association studies, in case of low dimension our method still performs well and can be an alternative to the perturbation method by Wu et al. (2010).

In our proposed methods, we only consider continuous responses. Extension to a binary response is natural and will be considered in our future investigation. Besides, our current methods were developed without prior knowledge. However, the kernel function actually allows for the inclusion of known information, such as the minor allele frequencies or association signals from an independent study. For example, weighted linear, quadratic, or IBS kernels can be constructed by assigning weights to variables individually. Thus, extension to weighted kernel is another direction that needs further investigation. With the next-generation sequencing data, identifying rare variants under the kernel machine framework has been a standard means in rare variants detection (Wu et al. 2011). Our method can also be applied to sequencing data under the KBT framework to improve power by integrating multiple kernel functions. This will also be investigated in our future work.

## Acknowledgments

This work was partially supported by grants from NSF (DMS-1209112 and DMS-1309156). Funding support for the GWA mapping: Maternal Metabolism-Birth Weight Interactions study was provided through the NIH Genes, Environment and Health Initiative [GEI] (U01HG004415). The datasets used for the analyses described in this manuscript were obtained from db-GaP at http://www.ncbi.nlm.nih.gov/sites/entrez?db=gap through dbGaP accession number phs000096.v4.p1. Code for implementing the method was written in R, and is available at https://github.com/hetao12/multi-kernel-test.

## Appendix

### Sketch Proof of Theorem 1

Under the null hypothesis, the leading order of the test statistic can be written as the sum of U-statistics of different orders. The asymptotic distribution can be then studied using U-Statistic theory (Lee, 1990). Under the local alternative, the test statistics can be decomposed into two parts, where the first part corresponds to the null distribution, and the leading order of second part converges to the location shift. See detailed proof in (He et. al 2018).

### Proof of Example 1

By the definition of centralized kernel in (3), we can obtain the centralized linear kernel as *K* (*x*_1_,*x*_2_) = (*x*_1_ — ***μ***)*^T^* (*x*_2_ — ***μ***), where ***μ*** = (*μ*_1_,*μ*_2_, …, *μ_p_*)*^T^* is the mean of random vectors **X**_*i*_, *i* = 1, …, *n*. Assuming the covariance matrix has decomposition **Σ** = ***Q^T^*****Λ*****Q*** with **Λ** being the diagonal matrix. Let 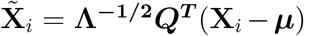, where it is obvious to see 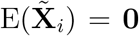 and 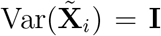, for *i* = 1, …, *n*. Noting that the centralized kernel can be written as

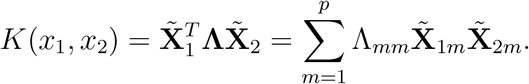

We can obtain our claim by letting 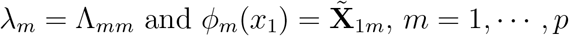.

### Proof of Example 2

We first derive the closed form of the centralized kernel function for the quadratic kernel 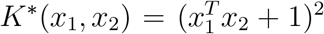. Decompose the kernel *K*^*^ into the sum of three parts

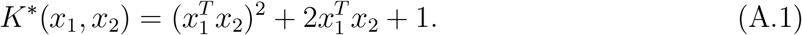

In the following we study each part separately, because the centralized function of the *K*^*^ is essentially the sum of individual centralized functions. For the constant 1, the corresponding centralized version is 0. Since we have studied the centralized version of inner product 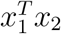 in Example 1, it remains to investigate the first term 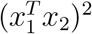. It is easy to show that

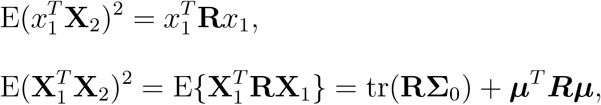

where **R** = (*R_ij_*) = **Σ**_0_ + ***μμ**^T^* is a constant matrix, and ***μ***, **Σ**_0_ are the mean and covariance matrix of **X**_*i*_ respectively. Thus the centralized version of 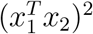 is

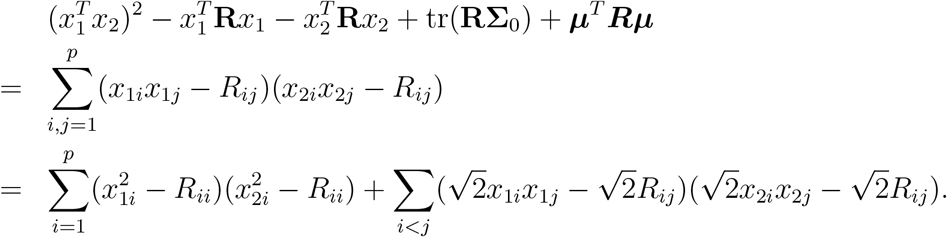

Combing the centralized expansions for the three terms in (A.1), we can rewrite p

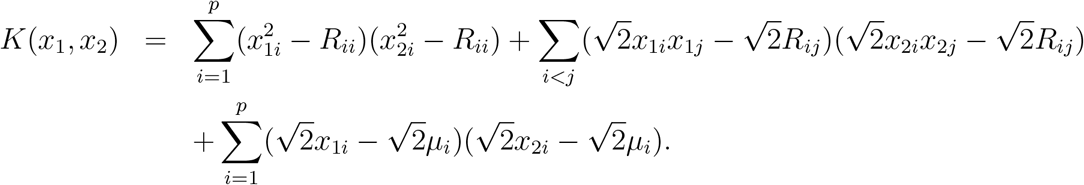

Assuming random vector 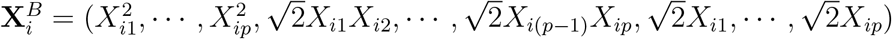 follow some distribution with covariance matrix **Σ** = ***Q**^T^* **Λ*****Q***, then we can achieve our conclusion, i.e., 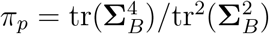, by performing the similar orthogonal transformations we proposed in the proof of Example 1.

### Proof of Example 3

For the IBS kernel taking the form of

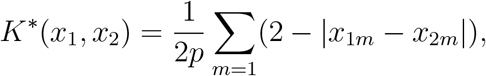

it is defied based on the total number of alleles shared identical by state (IBS) by two subjects at the SNPs within a SNP set. Noticing *X_im_* ∈ {0,1, 2}(1 ≤ *i* ≤ *n*, 1 ≤ *m* ≤ *p*), it is not difficult to verify that *K*^*^ has an alternative form of

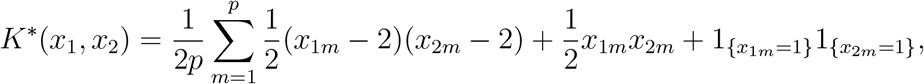

hence the centralized kernel has the following expansion

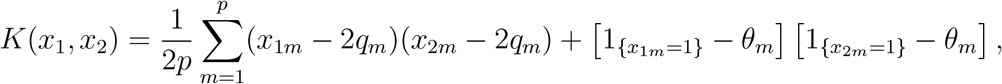

where *q_m_* is the minor allele frequency of the *m*th SNP, and *θ_m_* = P(*x_im_* = 1) = 2*q_m_*(1 - *q_m_*). Using the similar arguments as the proof of Example 1, we can obtain the result.

### Proof of Theorem 2

Assume condition (5) is satisfied for each candidate kernel *K_m_*, then

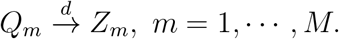

By using Cram*è*r-Wold device, 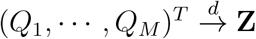 where **Z** follows a standard multivariate normal distribution. Then the first conclusion can be immediately obtained through the continuous mapping theorem. Since by Theorem 1 the power of single kernel test *Q_m_* depends on the location shift 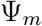, we thus denote the *m*^*^th kernel that has largest location shift 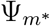 (i.e., 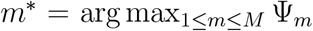) as the optimal kernel in the candidate set. Let 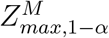 be the critical value of the maximum test at level *α*, then the power of the maximum test is

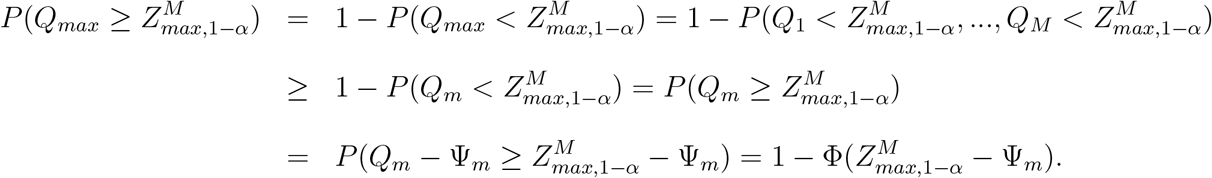

Therefore, the power of 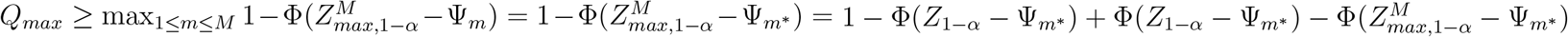, where we can reach the second conclusion because 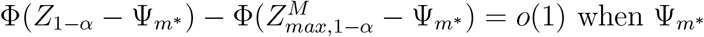 is large enough.

